# CD8+ T CELLS ASSOCIATE WITH FORMING GLANDULAR NODULES IN *TOXOPLASMA GONDII-*INDUCED PROSTATIC HYPERPLASIA AND HUMAN BPH

**DOI:** 10.64898/2026.08.17.745253

**Authors:** Tara D. Fuller, Rafael B. Polidoro, Doug W. Strand, Gustavo Arrizabalaga, Travis J. Jerde

## Abstract

**Background:** Chronic inflammation is the most common histological feature in Benign Prostatic Hyperplasia (BPH), and T cells are a key component of immune infiltrate. Advanced BPH is commonly associated with the formation of nodules, but it remains unclear whether a link exists among T cell infiltration, nodular development, and BPH progression. Using a *Toxoplasma gondii* (*T. gondii*) model and human specimens, we characterize the subtypes of T cells present during prostatic hyperplasia and their association with nodular development of the prostate.

**Methods:** Male CBA/j mice were intraperitoneally infected with *T. gondii* parasites, and flow cytometry was performed on the prostate to quantify the number of CD4+ and CD8+ T cells. Histology was used to score microglandular hyperplasia (MGH), and immunofluorescence was used to quantify and examine the locality of CD4+ and CD8+ T cells and compared that to human BPH tissue.

**Results:** We found that infecting male mice with *T. gondii* resulted in an increase of both CD4+ and CD8+ T cells in the prostate acutely and that CD8+ cells remained sustained at chronically. We also established the presence of glandular nodule formation at this timepoint through hematoxylin and eosin (H&E) staining. Immunofluorescence revealed that CD8+ cells were found proximal to forming glandular nodules relative to non-nodular glands. We also found more CD8+ cells localized to non-nodular glands in nodular BPH tissue versus non-nodular BPH tissue. Finally, we discovered a higher prevalence of CD8+ cells in *T. gondii* IgG+ patients than in IgG-patients. All *T. gondii* IgG+ patients exhibited nodular BPH, whereas all but one IgG-patient exhibited non-nodular BPH.

**Conclusions:** This study is the first to investigate the presence and location of CD4+ and CD8+ T cells within nodular and non-nodular BPH glands. We found an association of the presence of CD8+ T cells with nodular progression. This association held true in human prostate tissue. Translationally, CD8+ T cells may enhance nodular BPH progression, and *T. gondii* infection may promote this CD8+ T cell-mediated response.

## INTRODUCTION

Benign Prostatic Hyperplasia (BPH) is a progressive chronic condition characterized by a benign enlargement of the prostate and a collection of symptoms known as lower urinary tract symptoms (LUTS), an umbrella term that manifests in increased urinary frequency and urgency, nocturia, hesitancy, weakened stream, and pelvic pain [1]. BPH affects 50% of men by age 60 and 90% of men by the age of 90 and is commonly diagnosed through a combination of symptom scores, physical examination, ultrasounds, and blood tests to measure levels of prostate-specific antigen (PSA) [2, 3]. BPH has a global impact of 94 million cases in 2019 [4]. The estimated annual economic burden in the United States is $3.9 billion in treatment of symptoms alone [5]. Healthcare costs attributed to BPH include pharmaceuticals including α-blockers, phosphodiesterase-5 (PDE5) inhibitors, and 5α–reductase inhibitors (5-ARIs). In more severe cases, surgical removal of the prostate through transurethral resection (TURP) or holmium laser enucleation (HOLEP) may become necessary [6]. However, no treatment options currently exist that abrogate or reverse growth of hyperplastic glandular epithelium or fibromuscular stroma in BPH.

Prostates change histologically during BPH progression. Prostate tissue is comprised of epithelial, stromal, vascular, and ganglia histological compartments involving multiple cell types. Chronic inflammation during progressing disease results in cells secreting factors that promote a reactive microenvironment contributing to BPH progression. This highly reactive microenvironment is associated with the formation and progression of BPH nodules. Nodular BPH can be defined as stromal, glandular (epithelial), or mixed depending on the type of nodule(s) present. The most common type of nodular BPH is epithelial glandular [7]. Stromal nodules are often circumscribed, lack glandular features, and are smaller than glandular nodules, which are encapsulated and appear spongy from papillary infoldings. It is unclear how nodules are formed or progress, but developing nodular formations are associated with advancing symptoms, particularly inward growth eventually necessitating surgical intervention [7, 8]. Therefore, understanding mechanisms for nodular formation represents a key scientific gap.

Chronic inflammation is associated with developing BPH, and this microenvironment is heavily represented by T-cell infiltrate [9, 10]. T cells are known to associate with fibromuscular and glandular epithelial nodules in BPH [7, 9, 11]. T cells have been reported to be the largest proportion of gland-associated immune cells [12]. Therefore, it is possible that T-cell driven mechanisms could be involved in nodular formation and growth. However, the identity and quantity of these T cell subtypes are currently unknown. Furthermore, whether a difference in the composition of T cell subtypes exists between non-nodular and nodular BPH is also unknown, and understanding this may provide key mechanistic insights to BPH pathology.

No experimental models fully represent all features of BPH, and most work on nodule formation involves histological patterning by high dose steroid hormones. Our lab has developed a mouse model of human BPH using the unicellular parasite *Toxoplasma gondii (T. gondii),* which recapitulates key features of human BPH, including inflammation, reactive hyperplasia, voiding dysfunction, and notably, adenomatous/microglandular hyperplasia progressing to glandular nodule formation [1]. This model tying inflammation to formation of hyperplastic glandular nodular features proves to be highly useful in studying the mechanisms behind nodular formations. Acute *T. gondii* infection elicits a strong proinflammatory immune response characterized by the infiltration and activation of dendritic cells, monocytes, and T cells, all of which control *T. gondii* infection. However, inflammation becomes chronic as the parasites encyst and reactivate, allowing study of chronic inflammation-induced growth. Effector CD8+ T cells are one of the most critical sources of IFNγ and are required to combat *T. gondii* infection, while CD4+ T helper cells have been shown to play a crucial role in the maintenance of CD8+ T cell function [13, 14]. This type I immune response is crucial to controlling the infection and inducing conversion of the parasite into its latent form that can persist for the lifetime of the infected animal.

Previous studies from our lab show that T cells are the most expanded immune cell group in the chronic *T. gondii* model. However, the types of T cells that predominate in our *T. gondii* mouse model are currently unknown. The subtypes of T cells in BPH and their analogous correspondence to BPH features in the human prostate also remains unknown [15]. In this study, we characterize the number and locality of CD4+ and CD8+ T cells in relation to glandular nodules in our model. Furthermore, we present the translational impact of our work by showing the locality of these T cell subtypes in human nodular BPH. Finally, we show that a correlation exists between the number of CD8+ T cells in human nodular BPH and *T. gondii* seropositivity.

## MATERIALS AND METHODS

### Parasites and infections

*Toxoplasma gondii* parasites of strain PruΔhxprt + ldh2GFP (PruΔC32) were maintained as previously described [1, 16]. Tachyzoites were syringe-lysed three times with a 27 ½ gauge needle after parasites were approximately 70% egressed from the cells. Parasites were then washed twice, resuspended in 1X PBS, and counted with a hemacytometer.

Male CBA/j mice were housed in the Laboratory Animal Resource Center (LARC) at Indiana University School of Medicine under the National Institutes of Health Guide for the Care and Use of Laboratory Animals. When mice were 10-12 weeks old, they were intraperitoneally (i.p.) injected with either sterile 1X phosphate-buffered saline (PBS) or 40,000 PruΔC32 *T. gondii* tachyzoites and monitored daily for signs of infection. Surviving mice were then sacrificed at 14, 28, and 60 DPI (Supplemental Fig. 1).

### Tissue harvesting

Mouse prostates were bisected, and half of the prostate with attached urethra and bladder was fixed in 10% neutral buffered formalin (NBF) for 24 hours at 4° C or 48-72 hours at room temperature. The corresponding prostate half was removed from the urethra and bladder and minced in Dulbecco’s Modified Eagle’s Medium (DMEM) on ice to less than 1 mm^3^ prior to transfer in a 6-well plate containing a total of 5 mL of DMEM and 200 U/ml of type IV collagenase (cat. no. 17104019, Gibco). Samples were then placed in 37° C incubator with 5% CO_2_ for a total of 2-3 hours. A pipette was used to mix the samples every 15-30 minutes to break the cells apart. After incubation, cells and remaining tissue were filtered through a 70-micron filter and added to DMEM supplemented with 10% FBS, 1% antibiotic/antimycotic (gibco, ca. no. 15240062), and 2 mM EDTA (supplemented DMEM). Single cell suspensions were spun at 500g for 10 minutes at 4° C and processed further for flow cytometry.

### Flow cytometry

Single cell suspensions (∼5x10^5^ – 1x10^6^ cells) were suspended in a 1:100 dilution of TruStain FcX (anti-mouse CD16/32) antibody (ca. no. 101320, BioLegend) in FACS buffer for 10 minutes on ice to block nonspecific binding. To stain dead cells, samples were washed and resuspended in Zombie Aqua Fixable Viability Dye and 1X PBS (ca. no. 423102, BioLegend) at a 1:500 dilution for 20 minutes at 4° C. The following antibodies were used to detect cell surface receptors using a 2x mastermix of conjugated antibodies in FACS buffer at a 1:50 dilution for 20 minutes at 4° C: PerCP rat anti-mouse CD45 (ca. no. 561047, BD Pharmigen), PE/Dazzle 594 anti-mouse CD3 (ca. no. 100245, BioLegend), APC/Fire 750 anti-mouse CD8a (ca. no. 100765, BioLegend), and PE/Cyanine7 anti-mouse CD4 (ca. no. 100421, BioLegend). Unstained and fluorescence minus one (FMO) controls were used to set negative gates. After stain and wash, cells were fixed and permeabilized for one hour at 4° C with permeabilization buffer. The cells were then resuspended in FACS buffer, covered with aluminum foil, and placed at 4° C overnight. Flow cytometry analysis was performed at the Indiana University Simon Cancer Center Flow Cytometry Research Core Facility on the BD Biosciences LSR Fortessa.

UltraComp eBeads Plus Compensation Beads (cat. no. 01-3333-42, Invitrogen) were used for single color controls to minimize spectral overlap between fluorophores and establish appropriate voltages to record sample events. Unstained controls were also used to account for autofluorescence. Unstained and FMO controls were used to set negative gates, and 300,000 events were recorded for each sample. Samples were gated to exclude doublets using forward scatter width (FSC-W) by forward scatter area (FSC-A) and on cytoplasmic granularity using side scatter width (SSC-W) by side scatter area (SSC-A). Next, cellular debris was excluded by gating FSC-A by SSC-A. Zombie aqua fixable viability dye was used to gate out dead cells, followed by gating for immune cells (CD45), T cells (CD3), and T cell subtypes (CD8 and CD4). Samples were further analyzed using FlowJo software version 10.

### Histology processing and scoring

Bisected prostate tissue from organs used for flow cytometry were fixed overnight at 4° C in 10% neutral buffered formalin and further processed to heat-fixed paraffin-embedded tissue slides as previously described [1]. Slides were stained with hematoxylin and eosin (H&E) prior to visualization using a Leica DM 2500 microscope and DCF425C camera. Data acquisition was achieved using Leica Application Suite X 3.3.3.16958 software for mouse samples and Aperio ImageScope v12.4.6.5003 for human samples.

MGH scoring was achieved by capturing 20X images of forming glandular nodules at 1-3 levels of depth, defined as 100 microns between stained sections, per mouse. Images were blinded and scoring was achieved by averaging scores among the varying levels. Scoring was accomplished based on an established grading system [17, 18]. Briefly, focality was given a score of 0 (MGH is absent) to 4 (widespread throughout sample), and intensity was given a score of 0 (MGH is absent) to 3 (MGH in stroma, forming nodular structure). Focality and intensity were multiplied together to give a combined score per mouse, and overall score of each group was determined based on the averages of each group (control vs. infected).

### Immunofluorescence

#### Mouse prostate samples

Slides with tissue sections consecutive to H&E-stained slides were prepared as previously described [19]. The following primary antibodies were diluted in blocking solution and placed on the tissue slices overnight in the dark at 4° C: rat anti-CD45 at 1:800 (ca. no. NB100-77417, Novus Biologicals), rabbit anti-CD45 at 1:100 (ca. no. 70257S, Cell Signaling Technology), rat anti-CD4 at 1:60 (ca. no. 14-9766-82, Invitrogen), and rabbit anti-CD8a at 1:800 (ca. no. ab209775, Abcam). Secondary antibodies Alexa Fluor 568 or 647 anti-rat or anti-rabbit from Invitrogen were used to detect the primary antibodies, and nuclei were stained as stated before [1]. Finally, coverslips were applied using VECTASHIELD antifade mounting medium with DAPI (ca. no. H-1200-10, Vector Laboratories). Imaging was achieved using the Leica DM 6000 microscope, and data acquisition was achieved using Leica Application Suite X 3.3.3.16958 software.

#### Human prostate specimens

Human samples were obtained with full patient consent and according to the ethical standards of the Declaration of Helsinki at UT Southwestern Medical Center and placed in the UTSW biorepository (Author DWS). These samples were surgically collected through prostatectomy or resection from patients experiencing symptomatic BPH. Prostate tissue samples were matched with corresponding serum collection for *T. gondii* serum exposure positivity by assessing for IgG antibodies using the *Toxoplasma gondii* IgG ELISA (ENZYME LINKED IMMUNOSORBENT ASSAY) Immunoassay per manufacturer’s instructions (BIO-RAD, Hercules, CA, USA) and measured on a Biotek Synergy H1 Hybrid Reader. Specimens were paraffin-embedded, sectioned at 5 microns, and fixed to slides prior to staining with the following primary antibodies: mouse anti-CD45 at 1:100 dilution (ca. no. NBP1-79127, Novus Biologicals), rabbit anti-CD4 at 1:100 dilution (ca. no. ab133616, abcam), and rabbit anti-CD8 at 1:100 dilution (ca. no. ab93278, abcam). Primary antibodies were detected using anti-rabbit Alexa Fluor 488 or anti-mouse Alexa Fluor 568 or 647 conjugated fluorophores from Invitrogen as stated above. Specimen information is stored in the UTSW biorepository through Dr. Doug Strand’s laboratory: https://strandlab.net/research. Briefly, BPH prostate specimens in both BPH and Donor control groups averaged 64 and 62 years of age, respectively. BPH specimens were diagnosed by UTSW urologists and were obtained post Trans-urethral Resection of the Prostate (TURP) or complete prostatectomy. Tissues analyzed for this study were not currently treated with 5-alpha reductase (5ARI) therapy. Donor control groups were clinically negative for BPH symptoms and pathology analysis indicated no histological evidence of BPH. Both groups were confirmed to be negative for prostate cancer by pathology, both groups had similar body mass index numbers, and similar racial and ethnic make-up.

#### Analyses

Counting CD45+, CD4+ and CD8+ cells was achieved using ImageJ 1.53t software. Regions of interest (ROI) were drawn around the periphery of a forming glandular nodule or non-nodular gland and expanded outward to 100 microns. Cells were then counted manually using the cell counter plug-in. Criteria for determining CD4+ and CD8+ cells in mouse samples were colocalization of the CD4+ or CD8+ cell surface marker with the CD45+ cell surface marker.

To determine the total number of cells within each ROI, images of nuclear staining were cropped and re-opened in Ilastik software, where they were made binary using a training tool to threshold the images, followed by watershed to place natural breaks in the cells that were not completely separated. Total number of cells from the binary image were then counted in ImageJ through analyzing particles.

### Statistical analyses

Statistical analyses were performed using GraphPad Prism software 10.0.2. Two-way ANOVA was used to compare cell types between two groups and among multiple time points. When appropriate, p values were adjusted using Šidák correction. When evaluating cell types within the same sample, two-tailed paired student’s t test was used for column analysis and one-way ANOVA was used for group analysis among timepoints. To account for unequal sampling sizes, Mann-Whitney U test was used to analyze MGH scores between control and infected groups. All values were statistically significant when p ≤ 0.05, and sampling distribution was measured using standard error of the mean.

## RESULTS

### *T. gondii* infection drives leukocyte and T cell prostate tissue infiltration

To characterize T cell subtypes within the prostate, mouse prostates were assessed at 14, 28, and 60 DPI by flow cytometry (gating strategy – Supplemental Fig. 2). Our results show substantial leukocyte infiltration upon infection, as shown in CD45+ cells (Fig. 1a). Further analysis confirmed previous work from our lab demonstrating that the most abundant immune cell type is composed of T cells (Fig. 1b). We found that CD4+ T cell infiltrate was observed within two weeks of the infection (Fig. 2a), whereas CD8+ T cells remain sustained in the tissue for at least 60 days (Fig. 2b). Altogether, these results suggest that *T. gondii* infection can result in T cell infiltrate in prostate tissue, and that CD8+ T cells remain elevated in the tissue chronically in this model.

**Figure 1.**
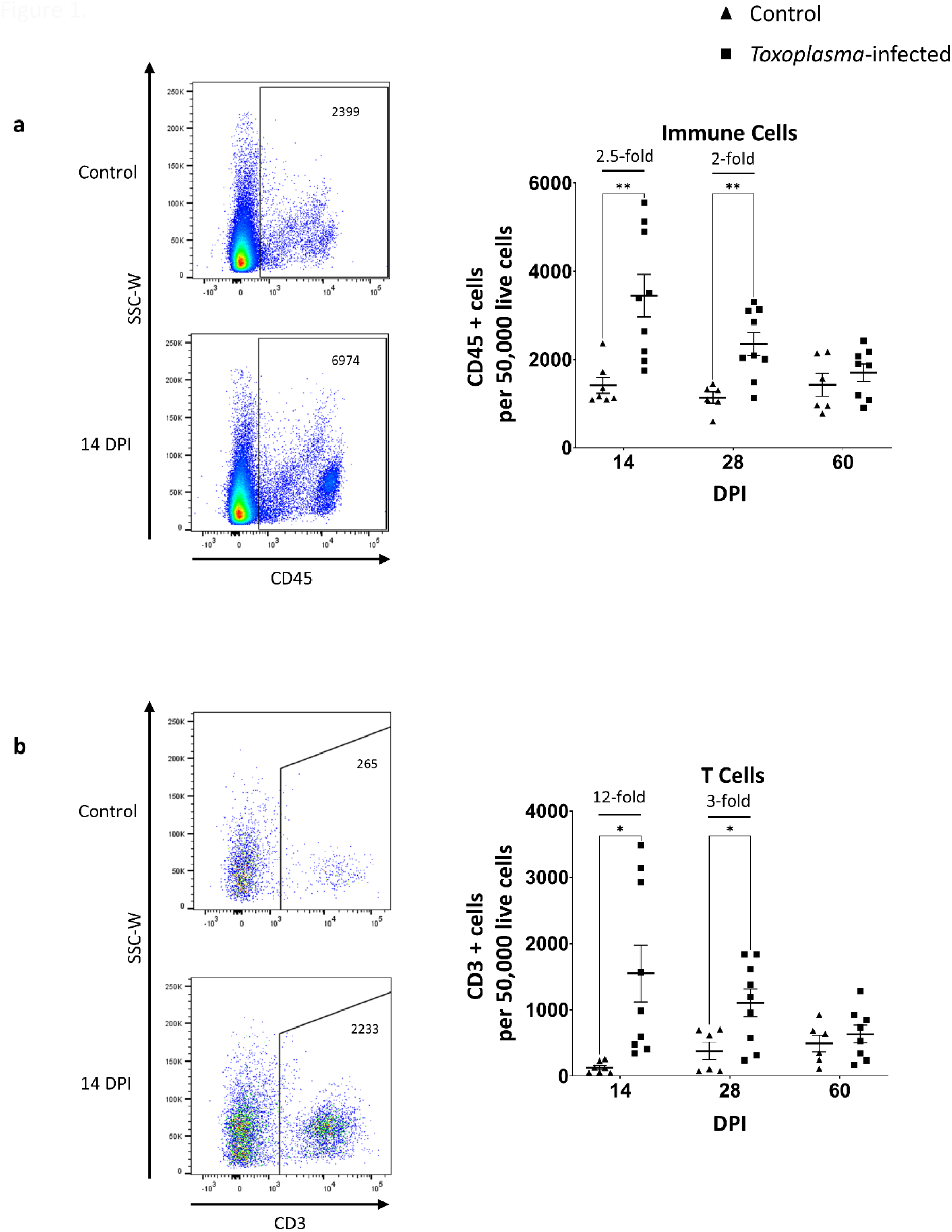
Leukocyte infiltrate in the mouse prostate is most pronounced during the acute phase of *T. gondii* infection. (a-b) (left) Gating strategy of live CD45+ and CD3+ cells for control and *Toxoplasma*-infected mouse prostates. (right) Flow cytometry was used to identify CD45+ and CD3+ cell populations, and numbers were normalized to 50,000 live cells at 14 (n = 7-9), 28 (n = 6-9), and 60 (n = 6-8) DPI from two separate experiments. Control prostate samples were compared to age matched infected mice. Data were analyzed using two-way ANOVA ± SEM followed by Šidák correction. *p value < 0.05, **p value < 0.005.

**Figure 2.**
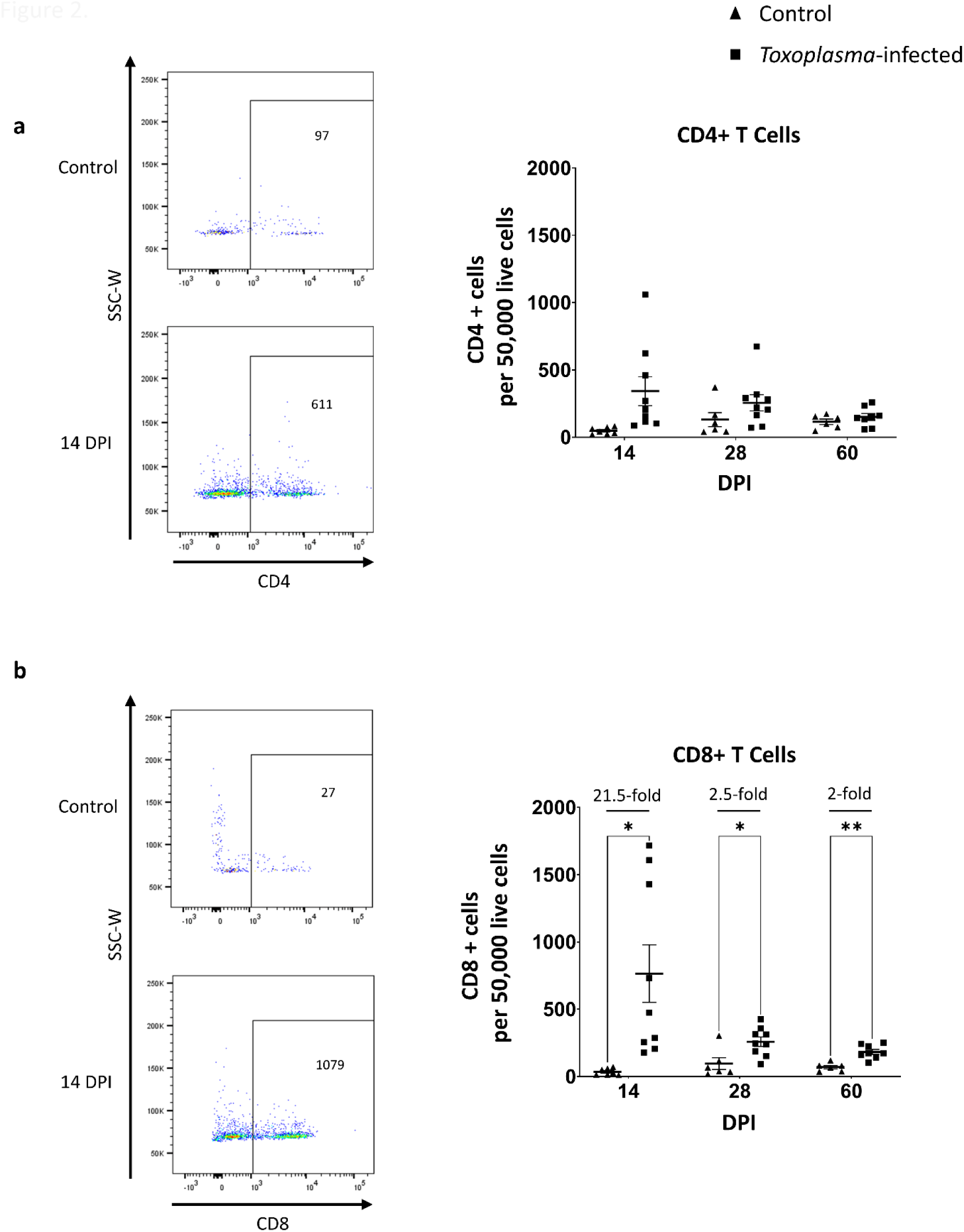
CD4+ and CD8+ T cells in the mouse prostate are the most pronounced early on in *Toxoplasma gondii* infection. (a-b) (left) Gating strategy of live CD4+ and CD8+ cells for control and *Toxoplasma*-infected mouse prostates. (right) Flow cytometry was used to identify CD4+ and CD8+ cell populations, and numbers were normalized to 50,000 live cells at 14 (n = 7-9), 28 (n = 6-9), and 60 (n = 6-8) DPI from two separate experiments. Control prostate samples were compared to infected. Data were analyzed using multiple unpaired t tests ± SEM followed by Holm-Sidak correction. *p value < 0.05, **p value < 0.005.

### Forming glandular nodules correlate with elevated T cell number in *T. gondii*-infected mouse prostates

To determine if glandular nodule formation corresponds temporally to induction of T cell subtypes, we assessed H&E stains on transverse prostate tissue sections from control and *T. gondii*-infected mice. Figure 3a shows hyperplasia consisting of mitotically active cells in 14 DPI prostates (right) in infected prostates. *T. gondii*-induced inflammation initiates a sustained microglandular hyperplasia (MGH) that forms nodular structures in chronic models (Fig. 3a) [1]. Here, we demonstrate that forming glandular nodules in hyperplasic prostates (criteria found in Supplementary Table 1) exhibit developing epithelial glandular nodules as early as 14 DPI (p = 0.0418) (Fig. 3b), corresponding to CD8+ cell infiltrate. These findings show that elevated T cell CD4+ and CD8+ subtype numbers are associated with increased reactive and microglandular hyperplasia.

**Figure 3.**
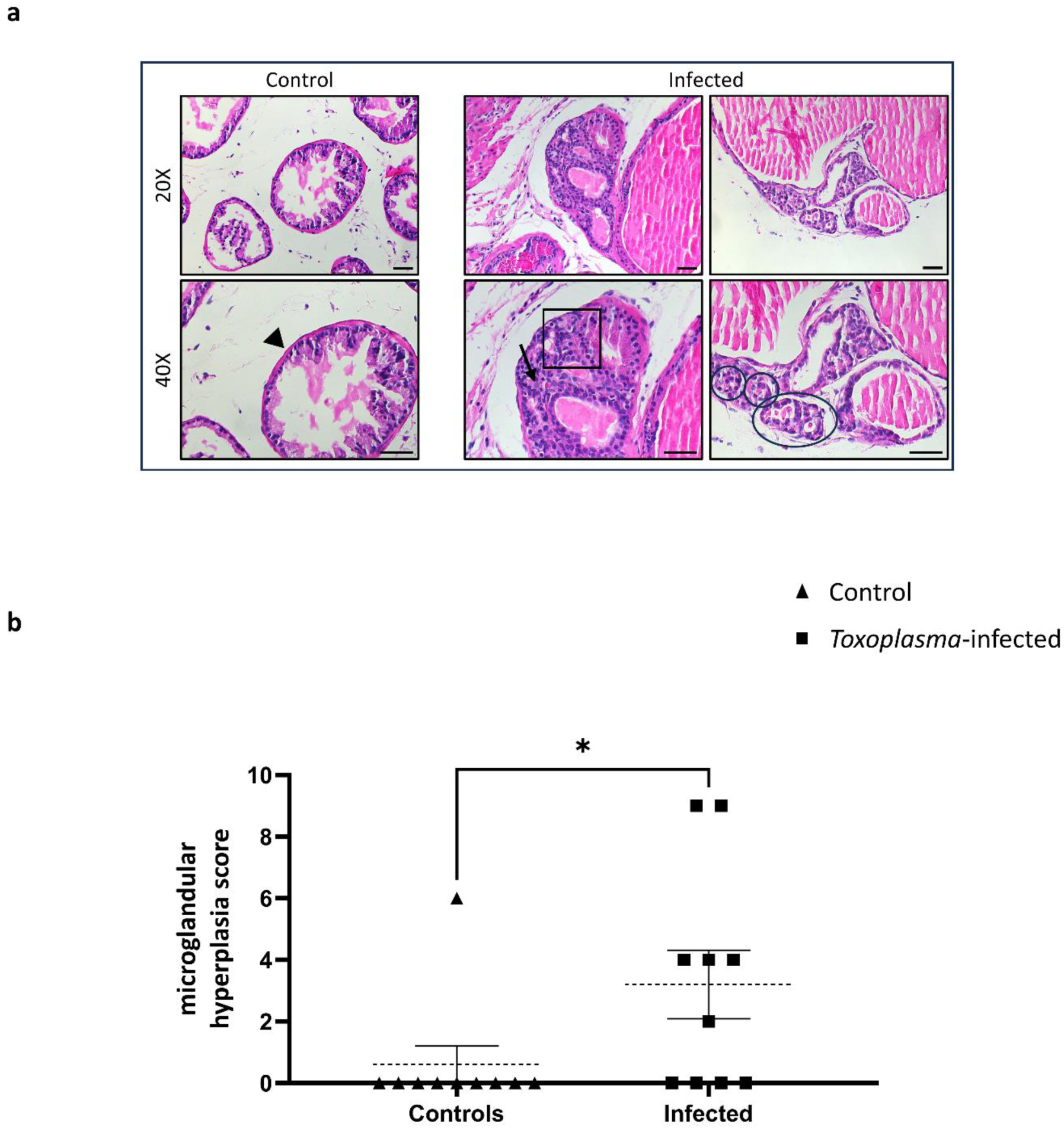
*Toxoplasma gondii* infection leads to early nodule formation in the mouse prostate. (a) Representative H&E images of control and 14 DPI mouse prostates at 20X and 40X objective. Black arrows reveal a single layer of pseudostratified columnar epithelium in control prostates (black arrowhead), while infected prostates exhibit hyperplasia (black box), actively dividing cells (black arrow), and small ring-like structures indicating forming nodules (black circles) as early as 14 DPI. (b) Microglandular hyperplasia scores from control and 14 DPI mouse prostates exhibiting forming nodules (n = 10, each group; 1 experiment) using the scoring system established in Table 1. *Toxoplasma gondii-*infected mouse prostates exhibit a higher microglandular hyperplasia score than controls. Data were analyzed using the Mann-Whitney U test. *p value < 0.05. Scale bars: 100 μm.

### CD8+ cells are consistently found within proximity of forming glandular nodules

To determine the locality of CD4+ and CD8+ cells, immunofluorescence identified CD4+ or CD8+ cells co-stained with the CD45+ immune cell marker (Fig. 4, 5) were quantified as localized within the boundary of, or within 100 microns of the periphery of glandular nodules or non-nodular glands. Data were represented per area (mm^2^) and per number (per 1,000 total cells). A consistently greater number of CD8+ cell subtypes were observed within or around glandular nodules compared to non-nodular glands within the same slice of tissue, (p = 0.0162; Fig 5). No difference in the number of CD4+ cells within or around both non-nodular glands or glandular nodules existed (Fig. 4). These results suggest that CD8+ T cells concentrate in close proximity to the forming glandular nodules and microglandular hyperplasia while CD4+ T cell infiltrate is homogeneous in the prostate tissue.

**Figure 4.**
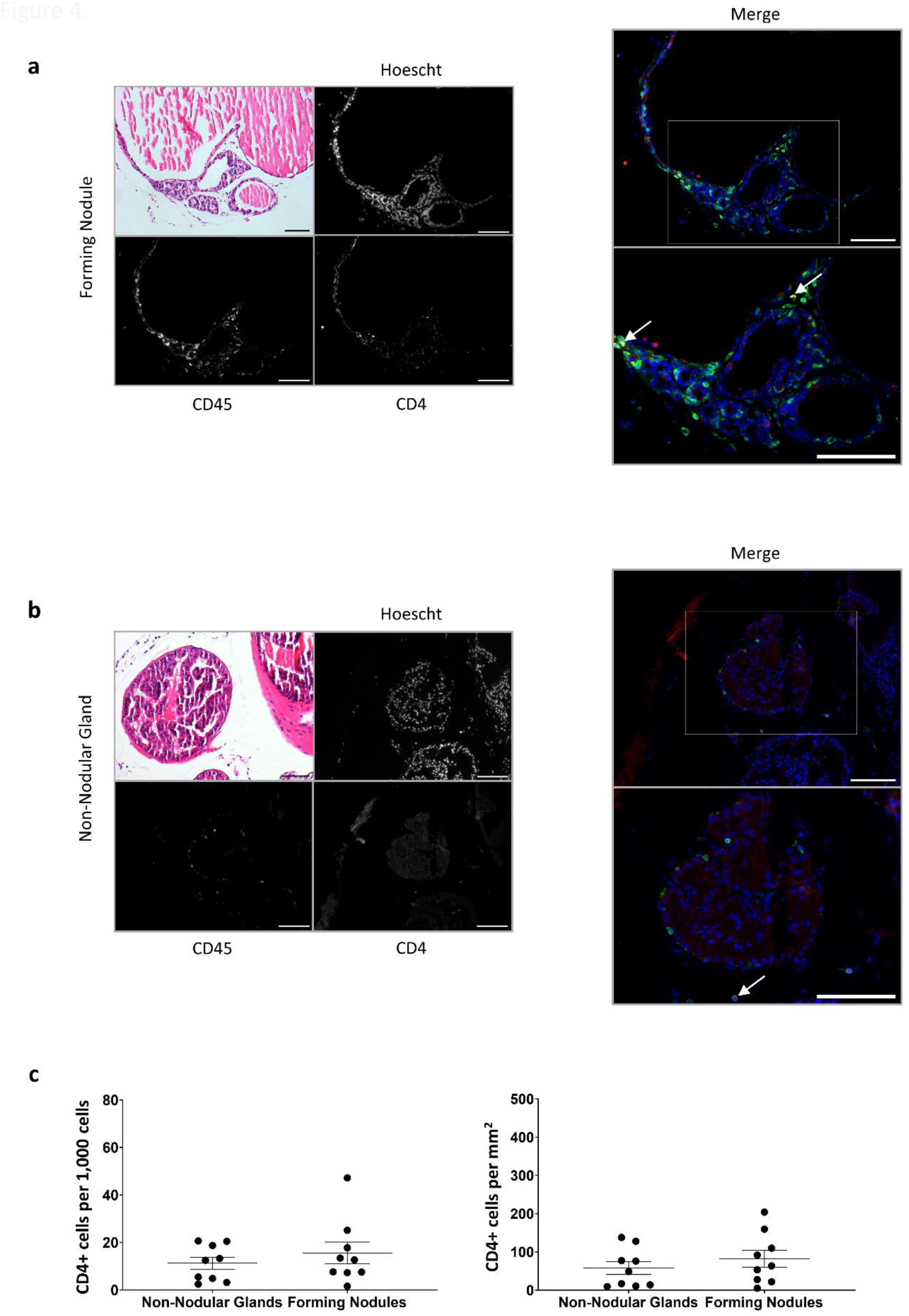
CD4+ T cells are dispersed in *Toxoplasma*-infected mouse prostates. (a, b) Representative image of immunofluorescence performed on 14 DPI (n=6) mouse prostates labeled with antibodies for CD45 (pan-immune cell marker) and CD4 (T helper cell marker). White arrows indicate colocalization (yellow) within 100 microns of forming nodule or non-nodular gland (c,d). Images taken with 20x objective from one experiment. (e) Quantification of T helper cells was achieved by counting CD4+ cells within 100 microns of forming nodules or non-nodular glands in 14 DPI mouse prostates using Fiji software. Results are expressed as both number of cells per area (mm2) and per 1,000 total cells and reveal a greater proportion of CD4+ T cells surrounding forming nodules as opposed to non-nodular glands. Data were analyzed using paired student’s t test ± SEM. n=9, 4 separate experiments. Scale bars = 100 μm

**Figure 5.**
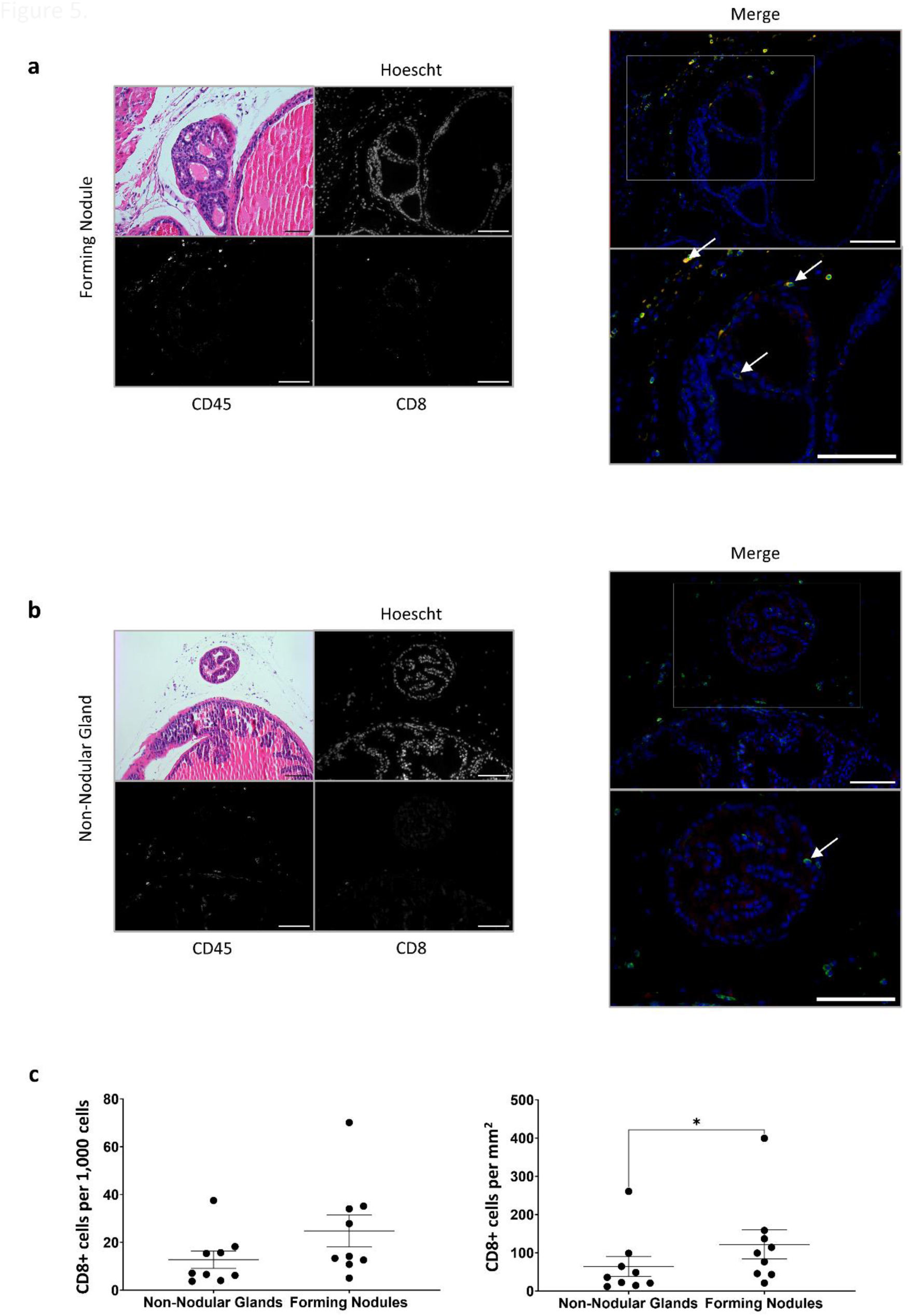
More CD8+ T cells than CD4+ T cells are proximal to forming nodules in *Toxoplasma*-infected mouse prostates. (a, b) Representative image of immunofluorescence performed on 14 DPI (n=6) mouse prostates labeled with antibodies for CD45 (pan-immune cell marker) and CD8 (cytotoxic T cell marker). White arrows indicate colocalization (yellow) within 100 microns of forming nodule or non-nodular gland (c,d). Images taken with 20x objective from one experiment. (e) Quantification of cytotoxic T cells was achieved by counting CD8+ cells within 100 microns of forming nodules or non-nodular glands in 14 DPI mouse prostates using Fiji software. Results are expressed as both number of cells per area (mm2) and per 1,000 total cells and reveal a greater proportion of CD8+ T cells surrounding forming nodules as opposed to non-nodular glands. Data were analyzed using paired student’s t test ± SEM. *p value < 0.05; n=9, 4 separate experiments. Scale bars = 100 μm

### Glandular nodular human BPH prostates exhibit higher numbers of CD8+ cells than non-glandular nodular BPH prostates

Human BPH specimens resected by TURP or complete prostatectomy were identified as exhibiting glandular nodule formations or not (Supplemental Fig. 3) and then further analyzed for immune cell subtypes. Due to the large and more complex nature of human BPH nodules, we evaluated these CD45+, CD4+, and CD8+ cells surrounding and within these nodules separately, as well as those surrounding non-nodular glands within the same tissue section (Supplemental Fig. 4, Fig. 6). Paired statistical analyses revealed no difference in CD4+ cells per 1,000 total cells or mm^2^ among glandular nodules, periglandular nodules or surrounding non-nodular glands. However, we found that more CD8+ cells localized to tissue overall within nodular BPH prostates than non-nodular BPH prostates per mm^2^ (p = 0.0021) (Fig. 7a). In addition, a significantly higher number of CD8+ cells were found in human prostate tissue from *T. gondii* IgG+ serum than IgG-tissue (p = 0.0031) (Fig. 7b), all five of which exhibited nodular growth, severe epithelial hyperplasia of a microglandular / adenomatous type. These results indicate that CD8+ cells are elevated in nodular BPH specimens and *T. gondii*-positive patients.

**Figure 6.**
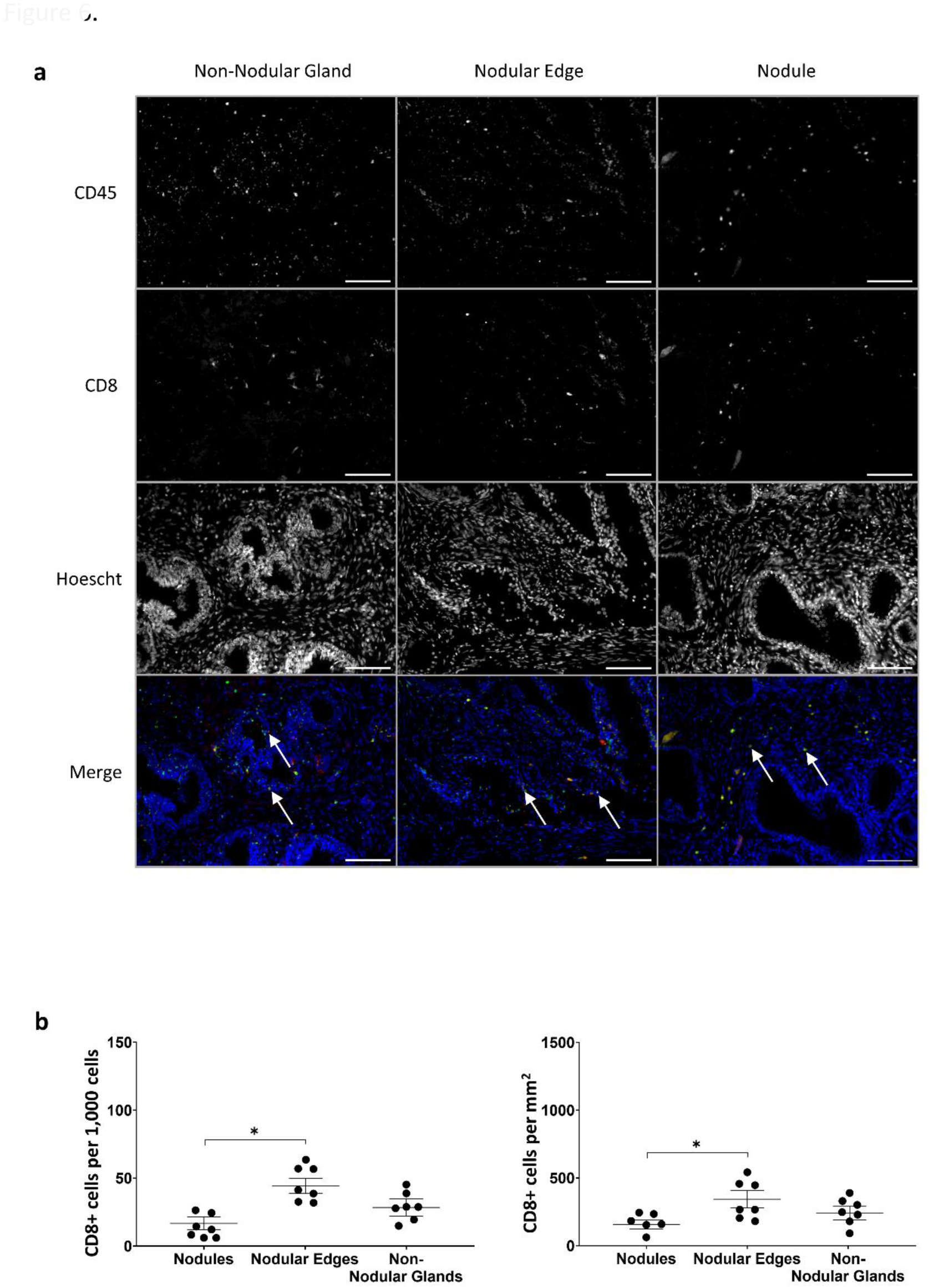
CD8+ T cells are found within and around nodules and non-nodular glands in comparable numbers. (a) Representative images of immunofluorescence performed on human prostate tissue exhibiting nodular BPH (n=4) labeled with antibodies for CD45 (pan-immune cell marker) and CD8 (cytotoxic T cell marker). White arrows indicate colocalization (yellow) within a nodule or 100 microns of a nodular edge or non-nodular gland. Images taken with 20x objective. (a) Quantification of CD8+ T cells was achieved as previously described. Results are expressed as both number of cells per 1,000 total cells and per area (mm2) and show no difference in CD8+ T cells within or around the nodule or non-nodular gland of each sample. Data were analyzed using paired student’s t test ± SEM with n=4. Scale bars = 100 μm

**Figure 7.**
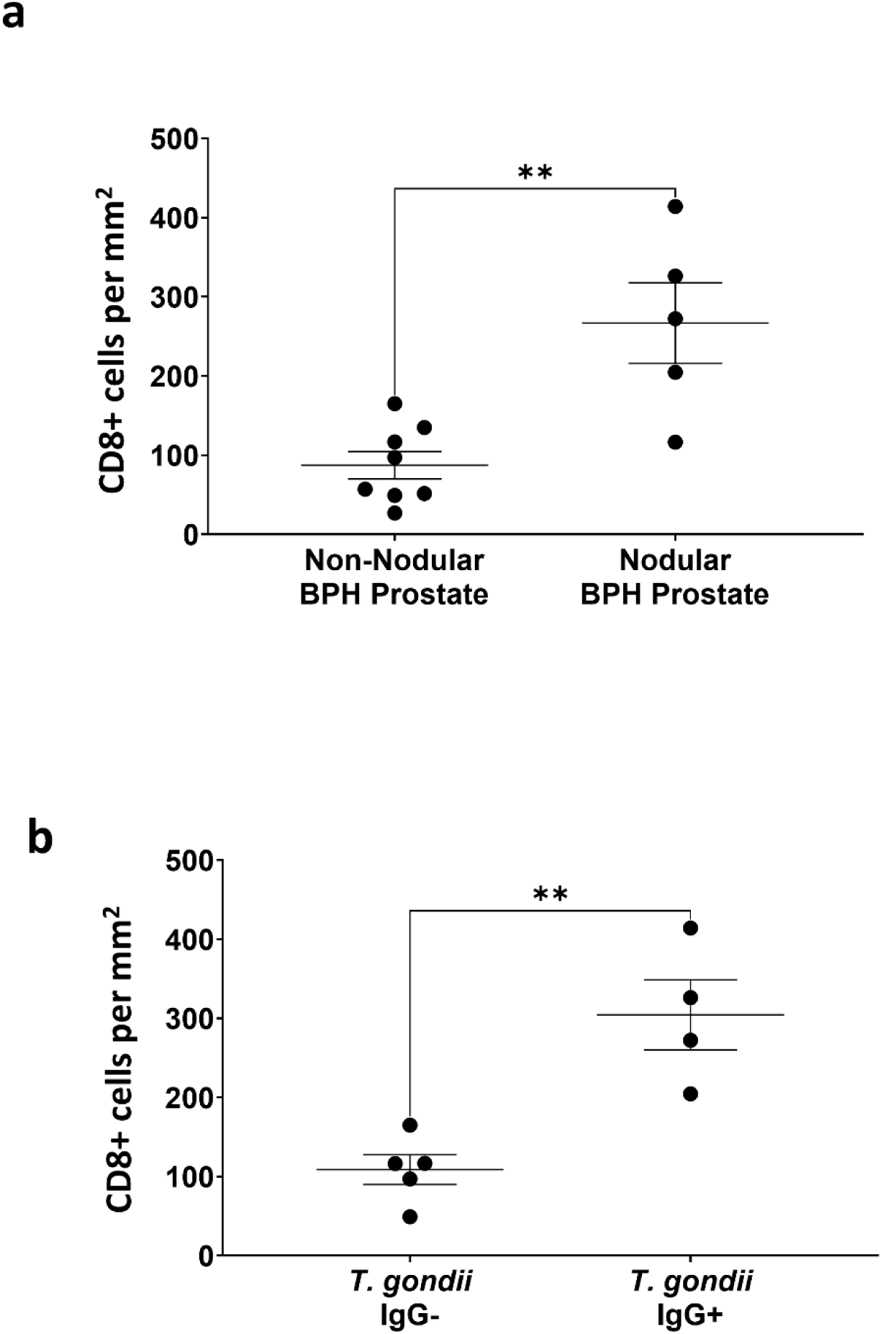
More CD8+ T cells are found in nodular than in non-nodular BPH tissue. (a) Quantification of CD8+ T cells was achieved as previously described. Results are expressed as both number of cells per 1,000 total cells and per area (mm^2^) (n=5-8). More CD8+ T cells exist within proximity of non-nodular glands in non-nodular BPH than around non-nodular glands in nodular BPH. In addition, patients who were *T. gondii* seropositive had more CD8+ T cells around non-nodular gland in their prostate tissue than those who were seronegative. Out of the 5 patients who were seronegative, 4 exhibited non-nodular BPH, whereas all 4 patients who were seropositive exhibited nodular BPH (b). Data were analyzed using unpaired student’s t test ± SEM. **p value < 0.005

## DISCUSSION

Using a mouse model of *T. gondii*-induced prostatic hyperplasia and glandular-like nodule formation combined with human BPH prostate specimens with glandular nodules, we found that CD8+ cells are specifically increased in glandular nodular prostates. In the mouse model, the CD8+ cells are concentrated around areas of microglandular hyperplasia and putative forming nodules, whereas CD4+ cells were more dispersed. Both T cell subtypes were expanded early in *T. gondii* infection during the acute phase, and CD8+ T cells remained elevated into chronic infection. In human prostates with glandular nodule BPH, CD8+ cells were high relative to non-nodular BPH though not localized inside the nodules per se. Interestingly, CD8+ cellular infiltrate was higher in the prostates of *T. gondii*-positive patients. In patients positive for *T. gondii* exposure, the parasite is known to infect the prostate, and studies suggest that parasite-manipulated cells home CD8+ T cells to areas of infection [20]. Future studies will be aimed at identifying the specific factors secreted from CD8+ T cells in the prostate and how they may be contributing to nodular formation.

The formation of epithelial glandular nodules exhibiting adenomatous or microglandular pathological structures is associated with advanced highly symptomatic BPH [7, 15]. Chronic and progressive inflammation also progresses with advancing BPH severity, but connections between inflammation and nodule formation remain unstudied. While T cell infiltrate is hallmark of progression of the disease, a clear localization of CD4+ and CD8+ T cells to glandular nodules and non-nodular glands has not been shown. The present study demonstrates that CD8+ cells localize to hyperplastic regions within the prostate, whereas CD4+ cells exhibit a more diverse distribution (Fig. 4, 5). CD8+ cells are higher in nodule harboring human prostates, particularly among those from patients testing positive for *T. gondii* exposure. These findings could be the result of multiple roles and mechanisms each T cell subtype plays [13, 14, 21]. For example, CD8+ cell subtypes secrete pro-proliferative and morphogenic cytokines such as IL-4, IL-17, and IL-6 in chronically inflamed tissue microenvironments [22–24]. In addition, CD8+ T cells generate IFNγ and TNFα, both of which can be involved in tissue damage and proliferation depending on the context [25, 26]. In the presence of inductive stroma, these cytokines promote the synthesis of growth factors and developmental morphogens that promote the proliferation of epithelial cells during repair and recovery and could drive the patterning of the epithelium into nodular structures in misdirected repair. A hypothesis generated from this work would be that glandular nodule formations could be an example of this type of tissue remodeling.

While the link between chronic inflammation and BPH progression in humans is well-established, recent research from our laboratory has shown an interesting clinical link between *T. gondii* exposure and the formation of advanced BPH in patients [18]. Stanczak et al. showed that patients BPH patients were significantly more likely to be previously exposed to *T. gondii* (as determined by positive serum IgG content) than age-matched donor patients histologically and clinically determined to be BPH free. *T. gondii*-positive BPH patients were statistically more likely to exhibit glandular nodules and microglandular / adenomatous hyperplasia that sero-negative pateints. In the present study, we showed that more CD8+ cells are present in *T. gondii* seropositive than seronegative BPH samples and that all seropositive BPH tissue is nodular in high CD8+ cell tissues. Further, *T. gondii* is known to promote a strong CD8+ T cell-mediated response. Collectively, these results hint at an association between parasitic infection, CD8+ cell infiltration, and BPH progression.

Although it remains to be known if *T. gondii* is a driver of BPH, experiments using therapeutics against CD8+ T cells could prove to be useful in treating disease progression. Pharmaceutical treatments that broadly inhibit CD8+ T cells remain to be addressed in the context of BPH. However, few drugs currently exist that directly target CD8+ T cells; the implications for side effects and lack of specificity remain problematic [27]. Studies in the past several years have shed light on the integral role that immune cells, particularly T cells, play in autoimmune diseases [26, 28, 29]. In fact, studies strongly suggest that BPH has an auto-immune component making the research of targeted anti-inflammatory drugs more crucial [26, 30, 31]. Pharmacological targeting toward T cell activation and cytokine production may prove to be a pivotal starting point for focused therapeutics. Future studies in our lab will be aimed at deciphering key mechanistic pathways induced by these secreted factors that contribute to epithelial cell proliferation and glandular patterning, as well as the subtypes of CD8+ T cells that are involved in our mouse model of *T. gondii*-induced prostatic hyperplasia and human BPH.

## ACKNOWLEDGEMENTS

The authors gratefully acknowledge Dr. Americo Lopez-Yglesias and the University of Texas Southwestern Medical Center Department of Urology Prostate Biorepository for their intellectual input, as well as Jerde lab member Hanyu Xia for his feedback and assistance with imaging analysis. This work was funded by National Institutes of Health-NIDDK (DK092366-01A1) and NIAID (DKI138255-01A1).

## AUTHOR CONTRIBUTIONS

Dr. Travis Jerde, Dr. Gustavo Arrizabalaga, Dr. Doug Strand, Dr. Rafael Polidoro, and Tara Fuller designed experiments. Dr. Jerde and Tara Fuller performed animal experiments, analyzed data and prostate pathology, and quantified staining. Dr. Jerde supervised staining quantification. Dr. Polidoro and Tara Fuller performed flow cytometry and analyzed data. Tara Fuller wrote the manuscript with assistance from Dr. Jerde and Dr. Arrizabalaga. All authors reviewed and edited the manuscript.

---

**Supplemental Figure 1.**
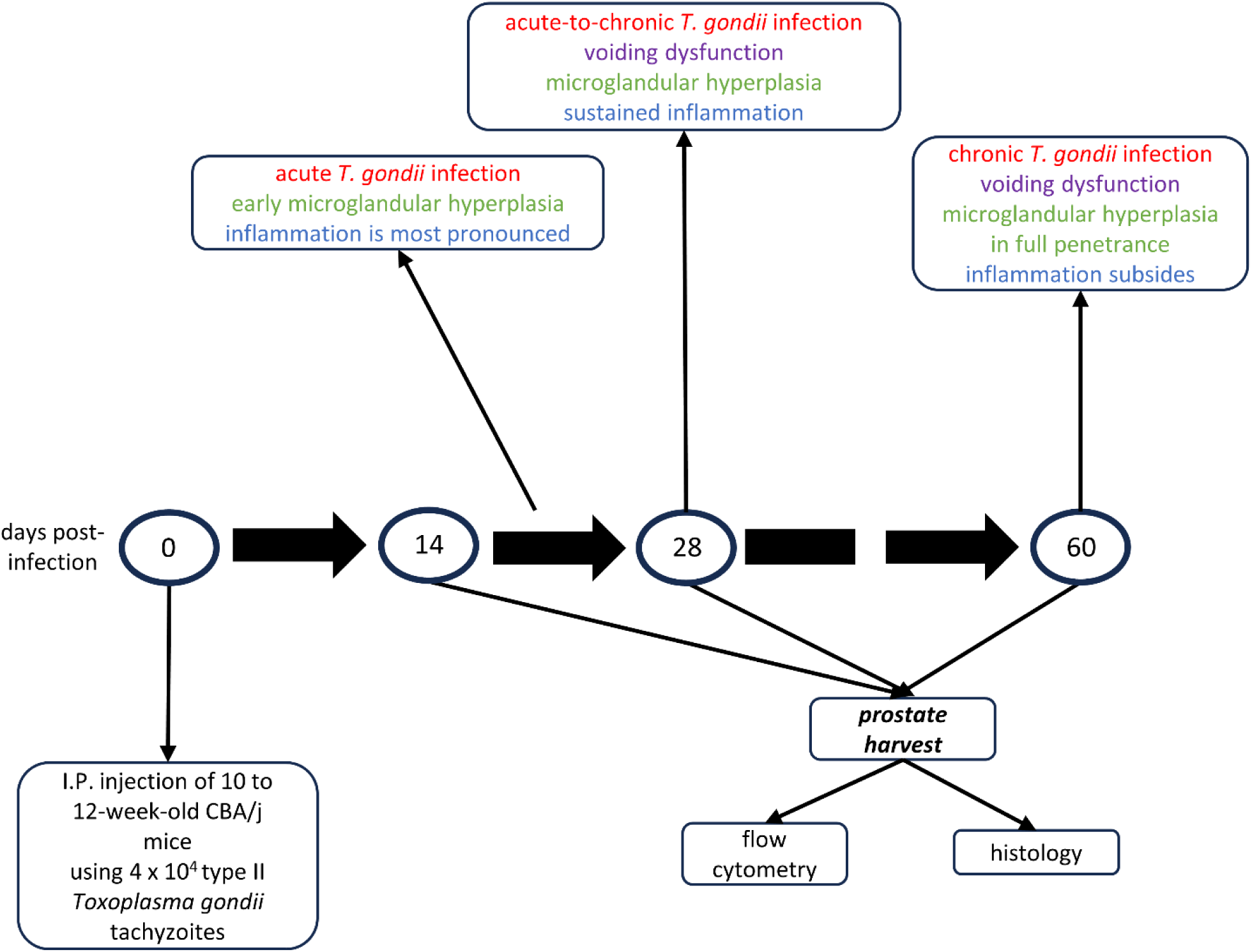
Schematic of experimental design. Ten- to twelve-week-old CBA/j mice were injected intraperitoneally with 40,000 PruΔC32 (PruΔhxgprt + ldh2GFP) *Toxoplasma gondii* tachyzoites or 1X phosphate buffered saline as a control. Prostates were harvested at 14, 28, and 60 DPI. Flow cytometry analysis was performed on half of each prostate; the other half was processed for histology. Each timepoint is characterized by pathological features that vary in severity and intensity (red = stages of infection; purple = symptoms; green = degree of microglandular hyperplasia; blue = level of inflammation).

**Supplemental Figure 2.**
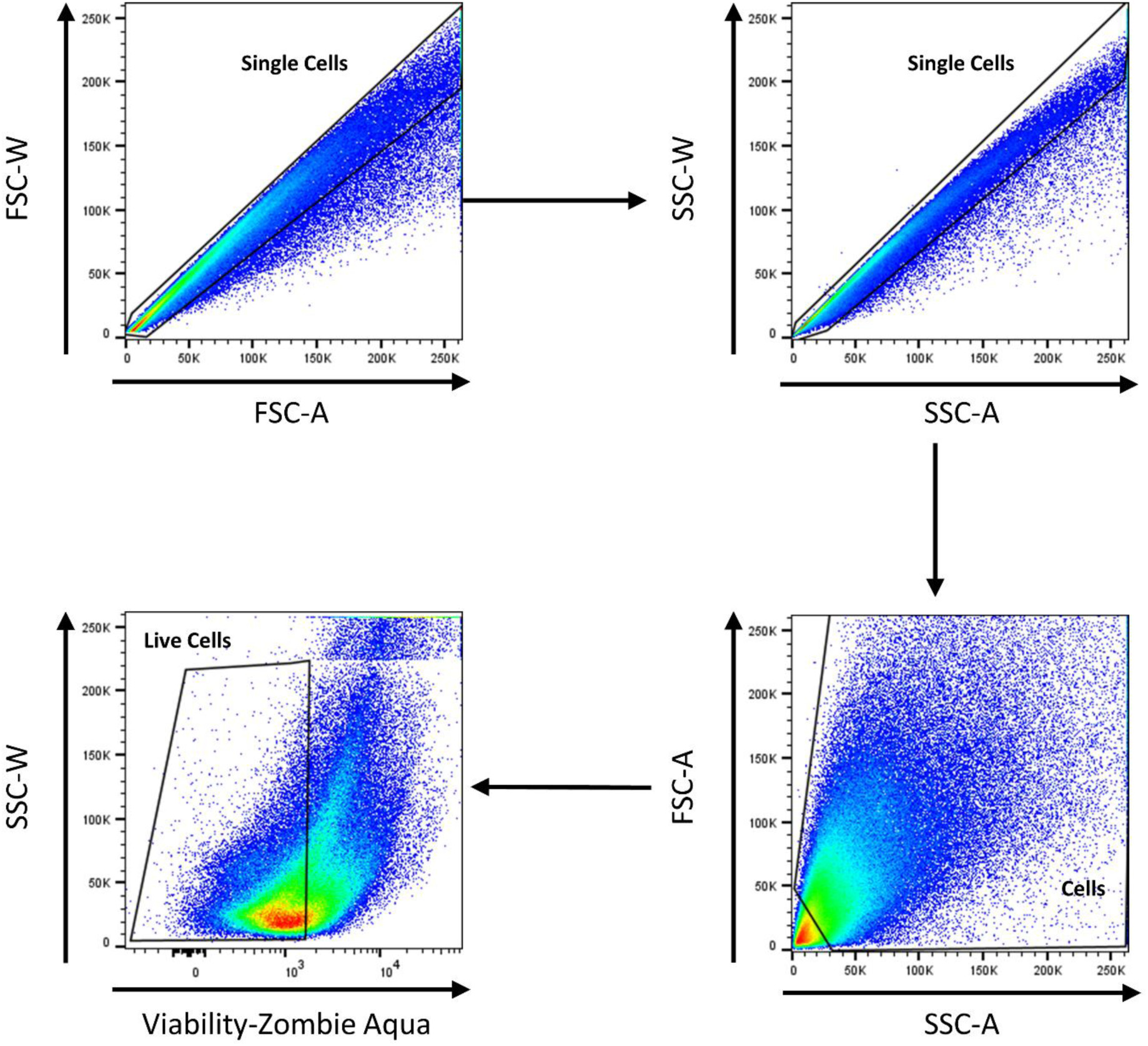
Flow cytometric gating strategy for CD4+ and CD8+ T cells in 14-day control and *Toxoplasma*-infected mouse prostates. Doublets were excluded from single cells using forward scatter followed by side scatter plots. Cellular debris and red blood cells were excluded from these single cells using forward by side scatter area. Prior to selection for CD45+ cell populations, zombie aqua viability stain was used to select for live cells based on the negative gating established by fluorescence minus one control.

**Supplemental Figure 3.**
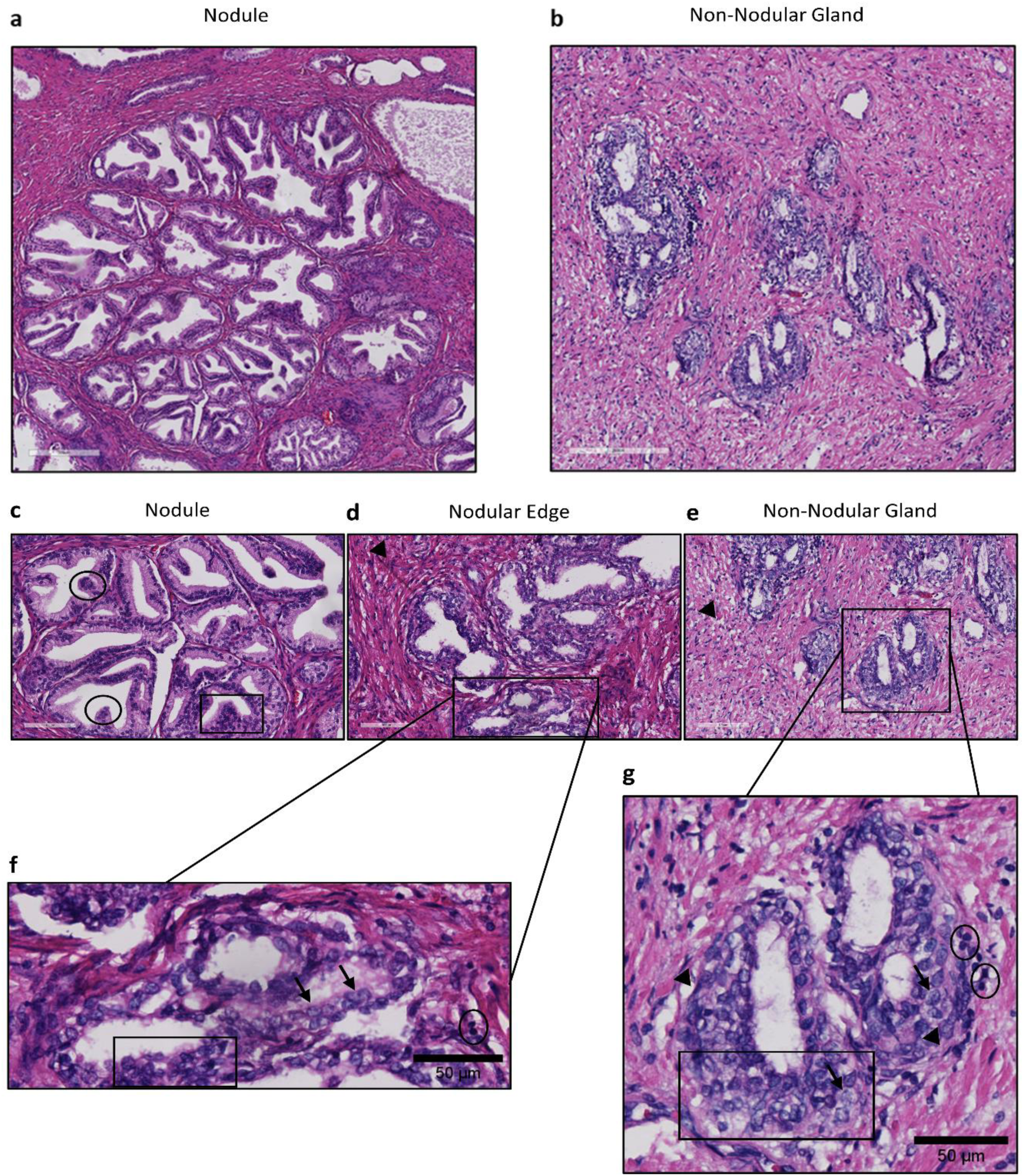
Similar features of prostate hyperplasia and nodular formation are seen in both *T. gondii*-infected mouse prostates and human nodular BPH. Representative H&E images of benign prostate hyperplasia (BPH) in human prostates showing features that are also seen in *T. gondii*–infected mouse prostates. (a) Nodular gland (scale bar: 300 mm) and (b) non-nodular gland (scale bar: 200 mm). (c) Epithelial nodule at 20X magnification that is characteristic of glandular BPH exhibits papillary infoldings protruding into the duct (black circles), as well as epithelial hyperplasia (black box) inside of the nodule. (d,e) The edge of the nodules show a thickened stroma (black arrowhead) and cribriform structures with a double-layered epithelium (black box) (d). Although (g) is considered to be a non-nodular gland, many cellular features of human BPH are seen in both non-nodular and nodular BPH (f) and (g). A magnified view shows prominent nucleoli in mitotically active cells and a clear cytoplasm (black arrows) that contribute to nuclear pleomorphism seen in epithelial hyperplasia (black box). Basal cell hyperplasia is also evident around the glands (black arrowheads), as well as inflammatory cells (black circles) juxtaposed to hyperplastic regions, undoubtedly contributing to a reactive microenvironment. Scale bars = 50 and 100 μm (c-g)

**Supplemental Figure 4.**
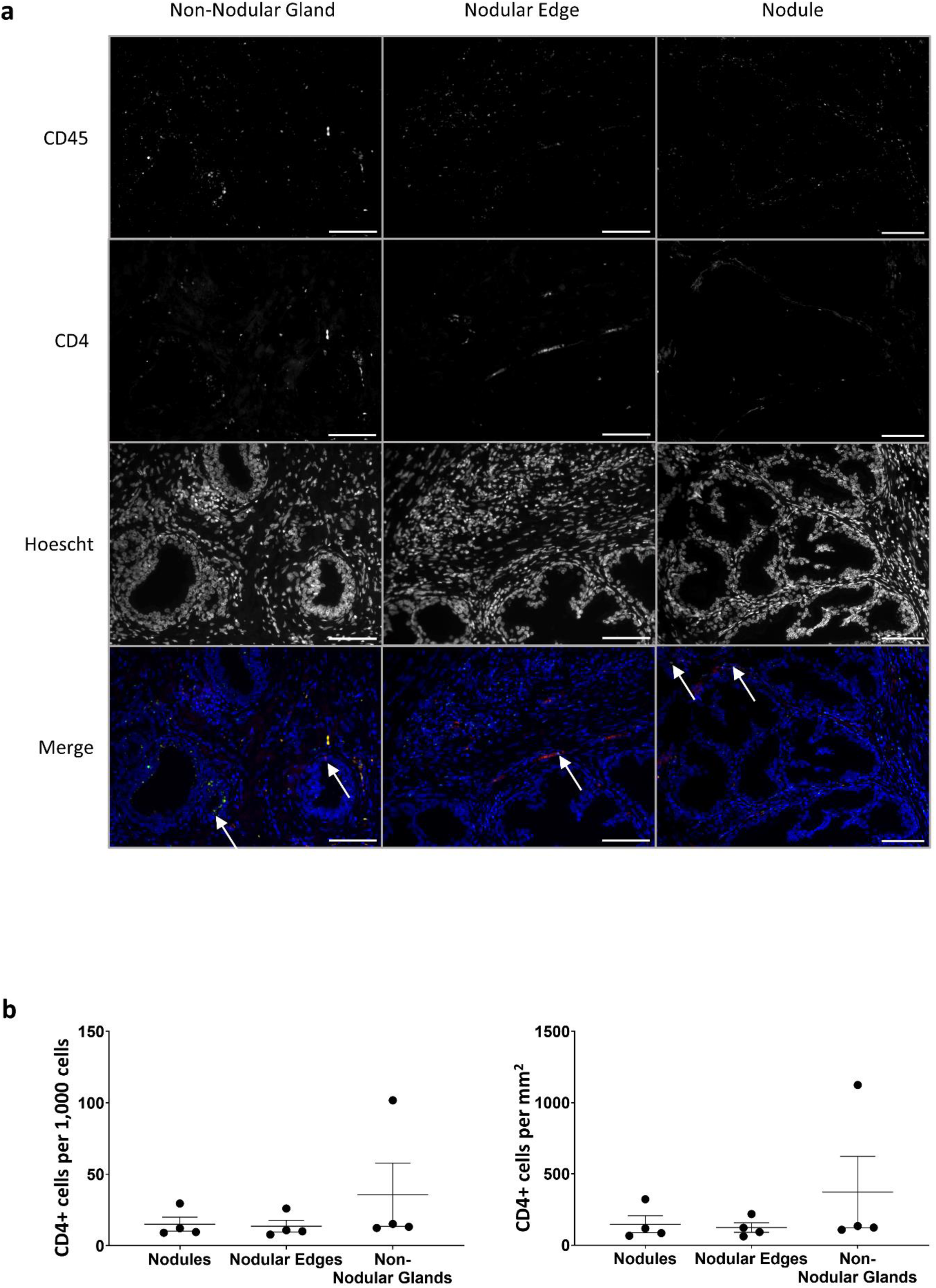
CD4+ T cells are dispersed in nodular BPH prostate tissue. (a) Representative images of immunofluorescence performed on human prostate tissue exhibiting nodular BPH (n=4) labeled with antibodies for CD45 (pan-immune cell marker) and CD4 (T helper cell marker). White arrows indicate colocalization (yellow) within a nodule or 100 microns of a nodular edge or non-nodular gland. Images taken with 20x objective. (b) Quantification of CD4+ T cells was achieved as previously described. Results are expressed as both number of cells per 1,000 total cells and per area (mm2) and show no difference in CD4+ T cells within or around the nodule or non-nodular gland of each sample. Data were analyzed using paired student’s t test ± SEM with n=4. Scale bars = 100 μm

