## Supplemental Information for "CD8+ T CELLS ASSOCIATE WITH FORMING GLANDULAR NODULES IN *TOXOPLASMA GONDII-*INDUCED PROSTATIC HYPERPLASIA AND HUMAN BPH"

Supplemental Figure 1.

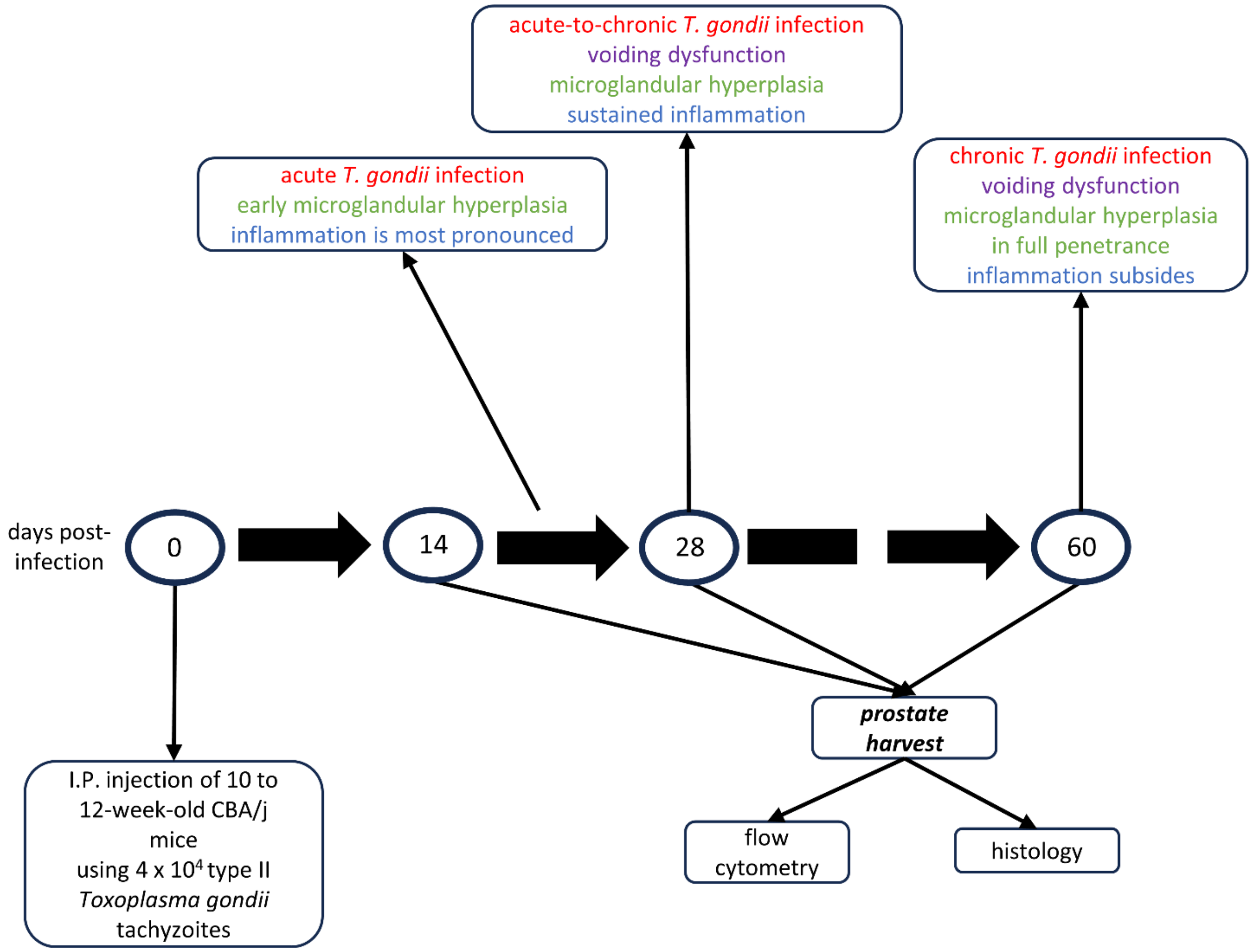


**Supplemental Figure 1. Schematic of experimental design.** Ten- to twelve-week-old CBA/j mice were injected intraperitoneally with 40,000 PruΔC32 (PruΔhxgprt + ldh2GFP) *Toxoplasma gondii* tachyzoites or 1X phosphate buffered saline as a control. Prostates were harvested at 14, 28, and 60 DPI. Flow cytometry analysis was performed on half of each prostate; the other half was processed for histology. Each timepoint is characterized by pathological features that vary in severity and intensity (red = stages of infection; purple = symptoms; green = degree of microglandular hyperplasia; blue = level of inflammation).

Supplemental Figure 2.

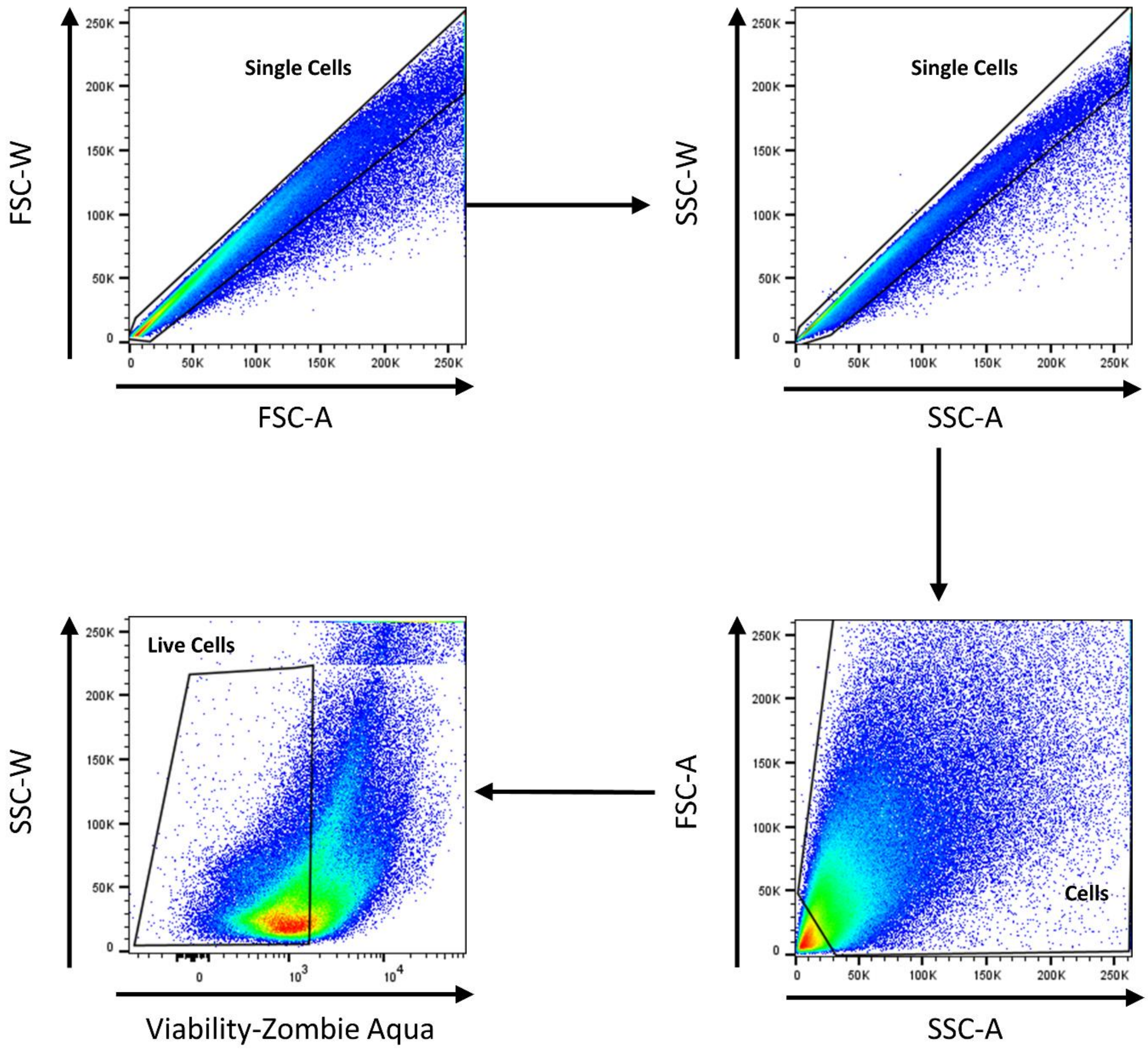


**Supplemental Figure 2. Flow cytometric gating strategy for CD4+ and CD8+ T cells in 14-day control and *Toxoplasma*-infected mouse prostates.** Doublets were excluded from single cells using forward scatter followed by side scatter plots. Cellular debris and red blood cells were excluded from these single cells using forward by side scatter area. Prior to selection for CD45+ cell populations, zombie aqua viability stain was used to select for live cells based on the negative gating established by fluorescence minus one control.

Supplemental Figure 3.

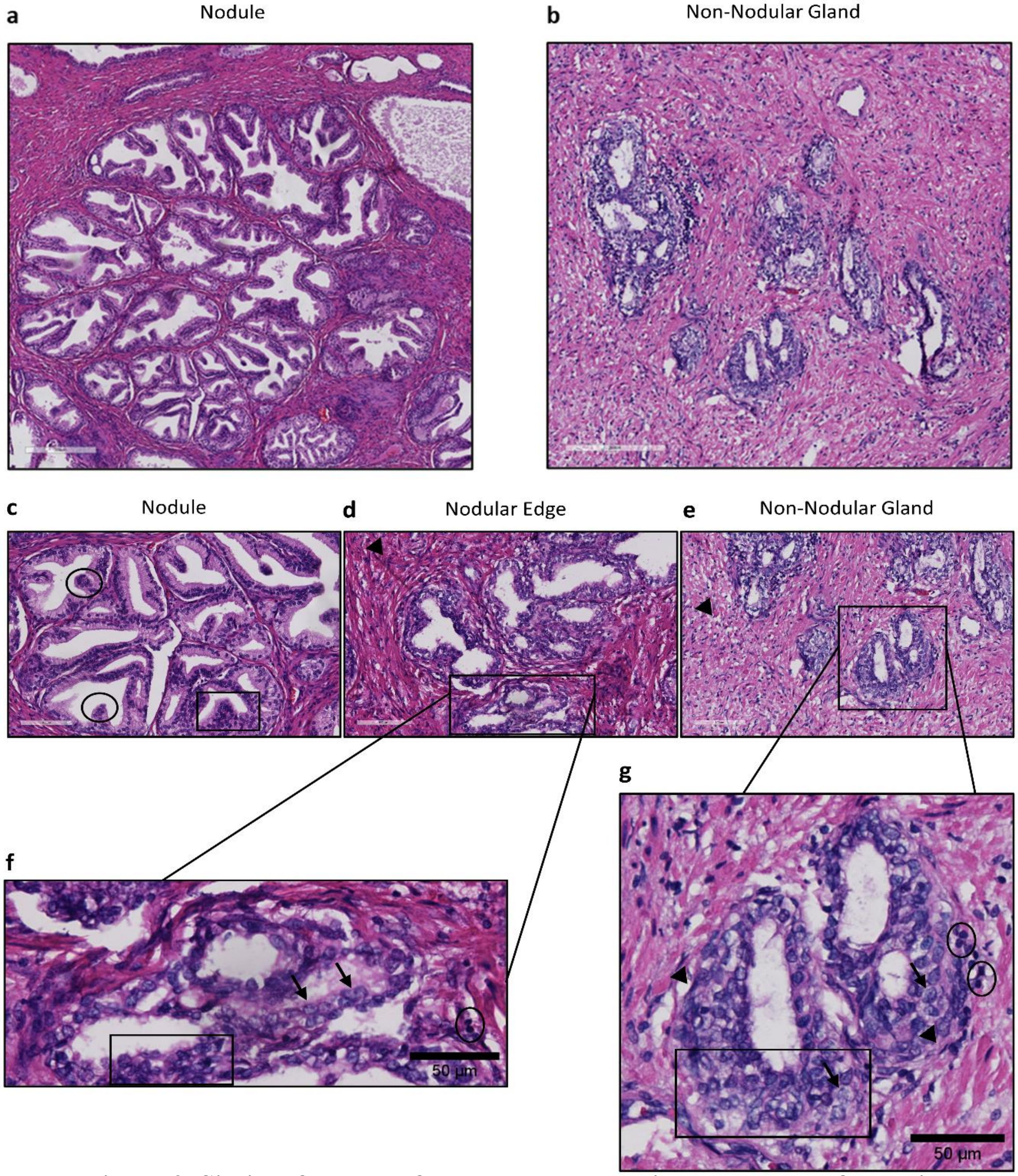


**Supplemental Figure 3. Similar features of prostate hyperplasia and nodular formation are seen in both *T. gondii*-infected mouse prostates and human nodular BPH.** Representative H&E images of benign prostate hyperplasia (BPH) in human prostates showing features that are also seen in *T. gondii*–infected mouse prostates. (a) Nodular gland (scale bar: 300 mm) and (b) non-nodular gland (scale bar: 200 mm). (c) Epithelial nodule at 20X magnification that is characteristic of glandular BPH exhibits papillary infoldings protruding into the duct (black circles), as well as epithelial hyperplasia (black box) inside of the nodule. (d,e) The edge of the nodules show a thickened stroma (black arrowhead) and cribriform structures with a double-layered epithelium (black box) (d). Although (g) is considered to be a non-nodular gland, many cellular features of human BPH are seen in both non-nodular and nodular BPH (f) and (g). A magnified view shows prominent nucleoli in mitotically active cells and a clear cytoplasm (black arrows) that contribute to nuclear pleomorphism seen in epithelial hyperplasia (black box). Basal cell hyperplasia is also evident around the glands (black arrowheads), as well as inflammatory cells (black circles) juxtaposed to hyperplastic regions, undoubtedly contributing to a reactive microenvironment. Scale bars = 50 and 100 μm (c-g)

Supplemental Figure 4.

**a**

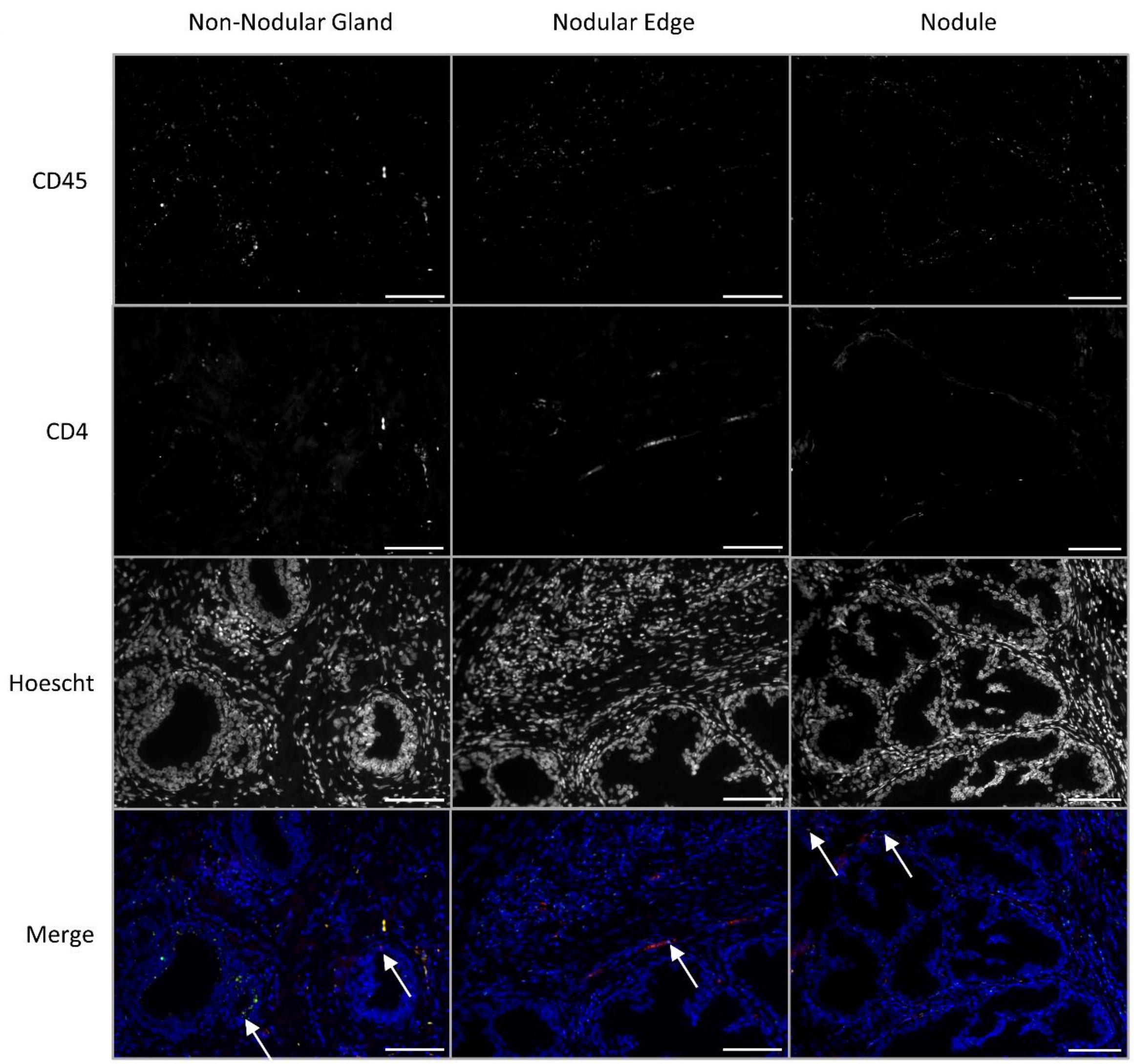


**b**

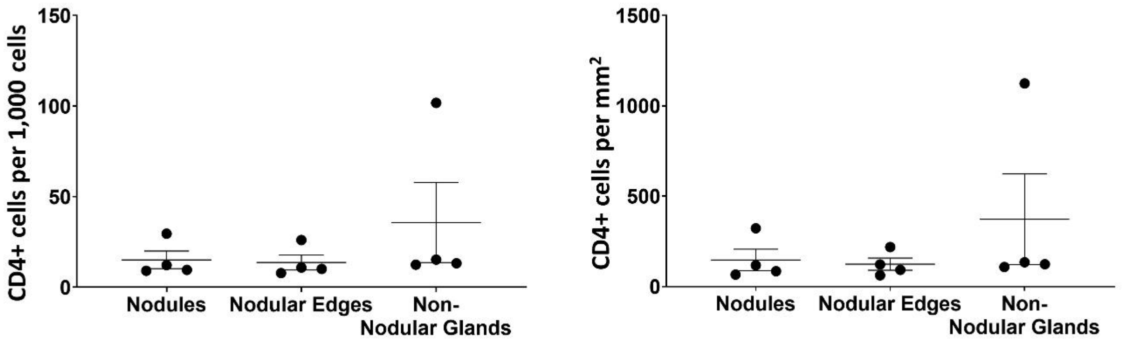

**Supplemental Figure 4. CD4+ T cells are dispersed in nodular BPH prostate tissue.** (a) Representative images of immunofluorescence performed on human prostate tissue exhibiting nodular BPH (n=4) labeled with antibodies for CD45 (pan-immune cell marker) and CD4 (T helper cell marker). White arrows indicate colocalization (yellow) within a nodule or 100 microns of a nodular edge or non-nodular gland. Images taken with 20x objective. (b) Quantification of CD4+ T cells was achieved as previously described. Results are expressed as both number of cells per 1,000 total cells and per area (mm2) and show no difference in CD4+ T cells within or around the nodule or non-nodular gland of each sample. Data were analyzed using paired student's t test ± SEM with n=4. Scale bars = 100 μm
